# Single-Molecule Peptide Fingerprinting

**DOI:** 10.1101/226712

**Authors:** Jetty van Ginkel, Mike Filius, Malwina Szczepaniak, Pawel Tulinski, Anne S. Meyer, Chirlmin Joo

## Abstract

Proteomic analyses provide essential information on molecular pathways of cellular systems and the state of a living organism. Mass spectrometry is currently the first choice for proteomic analysis. However, the requirement for a large amount of sample renders a small-scale proteomics study, such as single-cell analysis, challenging. Here we demonstrate a proof of concept of singlemolecule FRET-based protein fingerprinting. We harnessed the AAA+ protease ClpXP to scan peptides. By using donor fluorophore-labeled ClpP, we sequentially read out FRET signals from acceptor-labeled amino acids of peptides. The repurposed ClpXP exhibits uni-directional processing with high processivity and has the potential to detect low-abundance proteins. Our technique is a promising approach for sequencing protein substrates using a small amount of sample.

**SIGNIFICANCE:** Protein sequencing remains a challenge for small samples. A sensitive sequencing technology will create the opportunity for single-cell proteomics and real-time screening for on-site medical diagnostics. In order to resolve protein identity, we previously developed a computational algorithm that analyzes the ordered sequence of only two types of amino acids within a protein species. Through modification of a biological nanomachine, here we developed single-molecule fluorescence technology to linearize protein molecules and to read signals from labeled amino acids in an ordered manner. This proof of concept of singlemolecule fingerprinting will open a new door to single-molecule protein sequencing and pave the road towards the development of a new, fast, and reliable diagnostic tool.

## Introduction

Proteomic analyses provide essential information on molecular pathways of cellular systems and the state of a living organism (1). Thereby, for understanding of biological processes and their (dys)regulation, including disease, it is critical to monitor the protein composition of cells by sequencing (i.e. determination of the amino-acid sequence). Mass spectrometry is currently the first choice for protein sequencing. However, mass spectrometry analysis often fails to recognize minor species embedded among other dominant species since sequence prediction is made through analysis of complex spectral peaks (2). As many cellular proteins exist in low abundance (3), it is difficult to obtain large-scale proteomic information. DNA sequencing presents similar challenges, but they are overcome by amplifying DNA samples until a high signal-to-noise ratio is achieved. This solution cannot be applied to protein analysis since there is no natural machinery that can amplify proteins.

Single-molecule techniques have the potential to provide radically new protein sequencing tools that can quantify cellular proteins with accuracy as high as for mass spectrometry while requiring sample amounts as small as a single cell. However, despite several recent explorations (4–8), *bona fide* single-molecule protein sequencing has not yet been achieved due to the complexity that arises from primary protein sequences. Whereas DNA consists of only four building blocks (A, G, C, T), proteins are built from 20 distinct amino acids. Independent of the readout method of choice, full protein sequencing would require the detection of 20 distinguishable signals, which has so far not been demonstrated in single-molecule detection. Recently, our team and another have computationally demonstrated that read-out of only a subset of the 20 building blocks is sufficient to identify proteins at the single-molecule level (9, 10). In brief, the number of protein species in an organism is finite and predictable. Through bioinformatics-based comparison with proteomics databases, ordered detection of only two types of amino acids can still allow for protein identification. For example, ordered detection of cysteine and lysine residues, which can be modified using orthogonal chemistries, is sufficient to sequence the human proteome (10). We named this approach “single-molecule protein fingerprinting” to distinguish it from full protein sequencing. Here we demonstrate the first proof of concept of a single-molecule fingerprinting technology that reads out fluorescently labeled amino acids of synthetic peptides and a model cellular protein.

To obtain ordered determination of fluorescently labeled amino acids, we needed a molecular probe that can scan a peptide in a processive manner. We adopted a naturally existing molecular machinery, the AAA+ protease ClpXP from *Escherichia coli*. The ClpXP protein complex is an enzymatic motor that unfolds and degrades protein substrates. ClpX monomers form a homohexameric ring (ClpX_6_) that can exercise a large mechanical force to unfold proteins using ATP hydrolysis (11, 12). Through iterative rounds of force-generating power strokes, ClpX_6_ translocates substrates through the center of its ring in a processive manner (13, 14), with extensive promiscuity towards unnatural substrate modifications including fluorescent labels (15–17). Protein substrates are recognized by ClpX_6_ when they display specific disordered sequences such as the 11 amino-acid C-terminal ssrA tag (18). ClpX_6_ targets substrates for degradation by feeding them into ClpP_14_, a homotetradecameric protease that contains 14 cleavage sites and self-assembles into a barrel-shaped complex that encloses a central chamber (19).

## Results

### Single-molecule fingerprinting platform

To immobilize ClpXP (ClpX_6_P_14_) for single-molecule imaging, we biotinylated ClpX_6_ and bound ClpX_6_P_14_ to a PEG-coated quartz surface through biotin-streptavidin conjugation (Fig. 1a). A combination of total internal reflection fluorescence microscopy (TIRF) and Alternating Laser EXcitation (ALEX) imaging (20, 21) was used to monitor individual ClpXP complexes bind, translocate and degrade dye-labeled substrates in real time.

**Figure 1.**
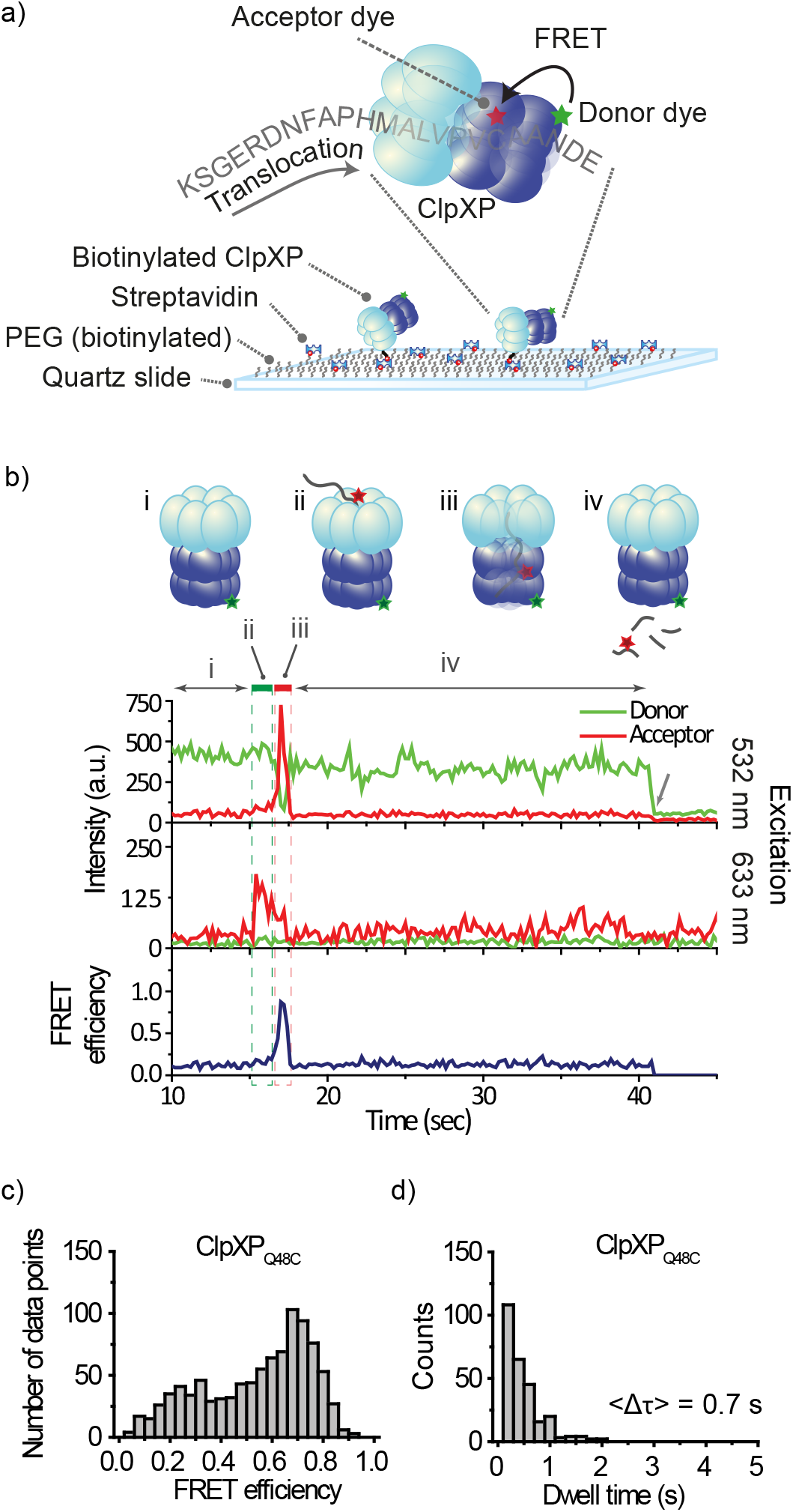
Single-molecule observation of ClpXP translocation. **(a)** Schematics of the single-molecule fingerprinting platform. Donor-labeled ClpXP is immobilized on a PEG-coated slide via biotin-streptavidin conjugation. ClpX_6_ recognizes an acceptor-labeled substrate (K-38-C_Cy5_-ssrA, **Supplementary Table 1**) and translocates it into the ClpP_14_ chamber upon which FRET occurs. **(b)** A typical fluorescence time trace. (i) The donor signal is from Cy3-labeled ClpXP (upper trace) upon green excitation (532 nm). (ii, green box) The sudden appearance of acceptor signal (time ~16 s) during acceptor-direct excitation with red (633 nm) reports on binding of acceptor-labeled substrate to ClpXP (middle trace). (iii, red box) The high FRET (time ~17 s) reports on the presence of the substrate in ClpP_14_ (top and bottom traces). (iv) Loss of fluorescence signal indicates the release of the substrate. The arrow at time ~40 s indicates the photobleaching of Cy3. (c) FRET distribution of stage iii. (d) Dwell time distribution of stage iii.

To detect the progression of fluorescently labeled amino acids through the ClpX_6_ pore with nanometer-scale accuracy, we employed FRET (Förster Resonance Energy Transfer) (22–24). We used two different types of model substrates for fingerprinting— short synthetic peptides and a small protein (the titin I27 domain). These substrates were labeled with acceptor fluorophores and were also appended with the ssrA tag. We constructed a FRET scanner by adding a fluorophore (donor) to the ClpP_14_ chamber (Fig. 1a).

We introduced cysteines to the Q48 residue of ClpP (ClpPQ_48C_), A139 (ClpP_A139C_) or F31 (ClpP_F31C_), labeled them with maleimide-functionalized fluorophores (**Supplementary Fig. 1a**), and evaluated the suitability for FRET-based substrate detection. ClpPQ_48C_ and ClpP_A139C_ showed higher FRET than ClpP_F31C_ (**Supplementary Fig. 1b**). Among the first two, ClpPQ_48C_ was chosen for our final scanner due to its higher efficiency of fluorophore labeling (see **Methods**).

The donor fluorophores on ClpPQ_48C_ are located near the center of the ClpP_14_ chamber, which is ~12 nm away from the substrate entry portal of ClpX_6_ (**Supplementary Fig. 1a**) (25, 26). This distance is longer than the Förster radius of a standard single-molecule FRET pair (~5 nm). This physical separation enables us to selectively detect signals from only the fluorophores (acceptors) on a protein substrate that have been translocated through a ClpX_6_ central channel. We obtained FRET time traces reporting on translocation, as shown in Figure 1b, by presenting a labeled peptide substrate to immobilized ClpXP complexes. The sudden appearance of acceptor signal during direct acceptor excitation indicates binding of acceptor-labeled peptide to ClpXP (Fig. 1b, **middle trace, stage ii**). The subsequent appearance of a high FRET state indicates translocation of the substrate by ClpX_6_ into the ClpP_14_ chamber (Fig. 1b, **stage iii**). When a slowly-hydrolyzable ATP analogue (ATP_*γ*_S) was used, the probability of high-FRET appearance decreased by one order of magnitude (**Supplementary Figure 2d**). Loss of FRET signal occurs upon the release of the dye-labeled peptide fragment (Fig. 1b, **stage iv**). When a cleavage inhibitor (DFP, diisopropyl fluorophosphate) was used (27) (**Supplementary Figure 2a-b**), the dwell time of high FRET increased 3.5-fold (**Supplementary Figure 2c**).

Our single-molecule fingerprinting concept requires detection of the order of fluorophores on a single substrate. To demonstrate fingerprinting, we functionalized a peptide with one type of fluorophore (Cy3) at the N-terminal site and a second type of fluorophore (Cy5) on an internal cysteine residue. We monitored the order in which the two fluorophores passed through Alexa488-labeled ClpP_14_ (Fig. 2a).

**Figure 2.**
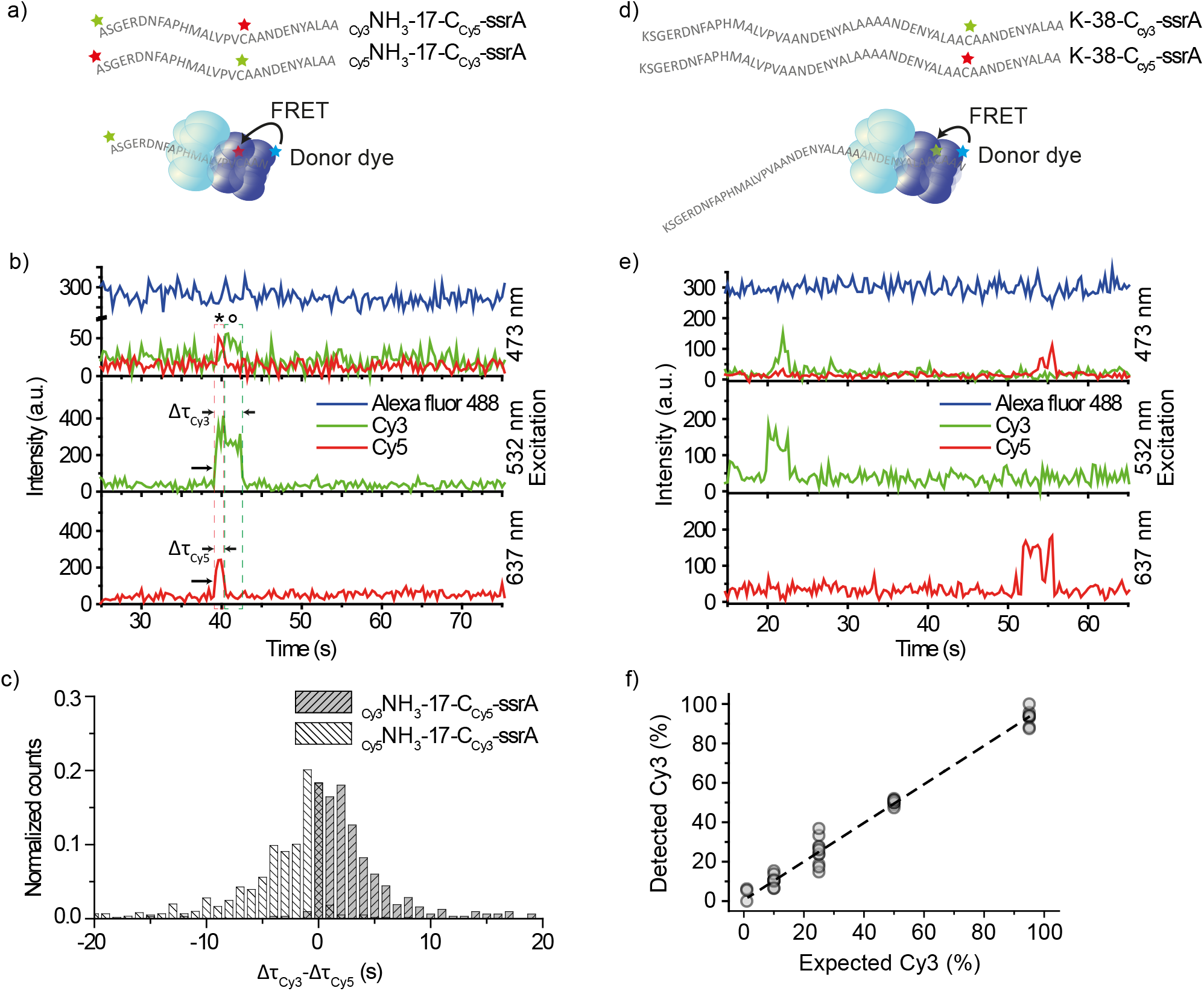
Single-molecule fingerprinting. **(a)** Substrates with two acceptor dyes were labeled at the N-terminal end and on cysteine residues (**Supplementary Table 1**). **(b)** A typical time trace for _Cy3_-NH_3_-17-C_Cy5_-ssrA from three-color ALEX (top) showed FRET between Alexa488 and Cy3; and Alexa488 and Cy5 upon excitation with blue laser light (473 nm). Concurrent signals from Cy3 (middle) and Cy5 (bottom) upon direct excitation, respectively with green (532 nm) and red (637 nm). For clarity, an arbitrary offset of 200 a.u. was applied to the Alexa488 trace, and the sum of the Cy3 and Cy5 signals was plotted (middle). For the original trace, see **Supplementary Figure 1e**. **(c)** Comparison of acceptor dwell times. The dwell time of Cy5 (Δτ_Cy5_) was subtracted from that of Cy3 (Δτ_Cy3_) for each event from _Cy3_-NH_3_-17-C_Cy5_-ssrA (grey). The mean of the distribution is 3.5 ± 4.90 (sec). In white is the same analysis for _Cy5_-NH_3_-17-C_Cy3_-ssrA. The mean of the distribution is −3.9 ± 5.67 (sec). **(d)** Substrates labeled with one acceptor dye. Cysteine residues of the substrates (K-38-C-ssrA) were labeled with either Cy3 or Cy5. **(e)** A time trace from substrates in [d]. At time ~20 s, a Cy3-labeled substrate binds (top, Alexa488-Cy3 FRET) and 532 nm direct excitation (middle). At time ~50 s, a Cy5-labeled substrate bind (top, Alexa488-Cy5 FRET) and 637 nm direct excitation (bottom). An offset of 200 a.u. applied to Alexa488. **(f)** The percentage of processed Cy3-labeled substrates was plotted against the expected percentage of Cy3-labeled substrates. The line is a linear fit (slope of 0.98 ± 0.02, intercept 0.58 ± 0.79, R^2^ = 0.99). Data points are from 100-s recordings, repeated ten times per condition (each *n* = 24.4 ± 1.50).

The positions of the Cy3 and Cy5 fluorophores relative to the ssrA tag on the substrate should dictate the order of Alexa488-Cy3 FRET and Alexa488-Cy5 FRET signals since an ssrA-tagged substrate is translocated through ClpX_6_ starting from its C-terminus. Figure 2b depicts a representative time trace obtained from a substrate (_Cy3_NH_3_-17-C_Cy5_-ssrA). The simultaneous appearance of Cy3 and Cy5 signals upon direct excitation with 532 nm and 637 nm (Figure 2b, **middle and bottom**, *t* ~ 40 s, indicated with arrows) indicates binding of a substrate containing both labels. In the FRET trace (Figure 2b, **top**), Alexa488-Cy5 FRET (marked * in the time trace) was observed before Alexa488-Cy3 FRET (marked °). This observation confirms that the ClpXP fingerprinter reads an ssrA-tagged substrate from the C-terminal to the N-terminal site.

We applied this fingerprinting scheme to the titin I27 domain. We labeled two Cys residues of titin (Cys64 and Cys80) with acceptor fluorophores. Because we did not have control over which dyes were attached to which Cys residues, we tagged both residues with the same dye, Cy5. Using Cy3 as a donor, we observed two separate FRET peaks within the time trajectories (**Supplemental Figure 3a**). The time interval between the two peaks was elongated when ATP_*γ*_S was mixed with ATP (**Supplemental Figure 3b**), indicating that the two peaks represented the sequential probing of Cys80 and Cys64 residues.

We extracted the length of time that Cy3 and Cy5 acceptor fluorophores were engaged with ClpXP (Δτ_Cy3_, Δτ_Cy5_). We observed positive differences in dwell time (Δτ_Cy3-Cy5_ = Δτ_Cy3_ – Δτ_Cy5_, < Δτ _Cy3-Cy5_> = 3.5 sec) for a substrate with N-terminal Cy3-labeling and internal Cy5-labeling (Fig. 2c, **grey**, _Cy3_NH_3_-17-C_Cy5_-ssrA). For a substrate with exchanged dye positions (Cy5NH3-17-CCy3-ssrA), we observed negative differences (Fig. 2c, **white**, < Δτ_Cy5-Cy3_> = −3.9 sec). Thus, dye-labeled amino acids located closer to the C-terminal ssrA tag were retained in the ClpXP complex for shorter amounts of time than labeled amino acids located more closely to the N-terminus. We can conclude that our fingerprinter can detect dyes in an order matching the amino-acid sequence. The ordered disappearance of the Cy3 and Cy5 signals further implies that uncleaved or partially cleaved substrate does not accumulate within the ClpP_14_ chamber, which would otherwise hamper accurate fingerprinting.

### Performance of the single-molecule fingerprinter

A single-molecule fingerprinter should perform without any bias to fluorophores and with high dynamic range. To determine the sensitivity of our fingerprinter, we performed a population study in which ClpP_14_ was labeled with donor fluorophore (Alexa488) and substrate peptides were singly labeled with either Cy3 or Cy5 as an acceptor fluorophore (Fig. 2d–e). We mixed Cy3- and Cy5-labeled substrates in varying proportions (1:99, 10:90, 25:75, 50:50, 95:5) and quantified the number of translocation events. We observed a linear relationship between the percentage of Cy3-labeled substrates we detected versus the expectation, with an offset of 0.58 ± 0.79 % and a slope of 0.98 ± 0.02 (adjusted R^2^ = 0.99) (Fig. 2f). We conclude that both FRET pairs are detected with equal sensitivity, and that our FRET scanner has the potential to detect low abundance proteins.

Our previous computational analysis indicated that the precision of our fingerprinting method would be enhanced if the distance between labeled cysteine and lysine residues could additionally be determined as well as their order (10). A uniform speed of the scanner, represented by ClpX_6_, is crucial to extract distance information. To determine whether the processing time of ClpXP is proportional to the length of protein substrates, we determined the processing times (the dwell time of fluorescence signals emitted by Cy5 labels on substrates, upon direct excitation) for three peptides (29, 40, and 51 amino acids (AA) in length; see **Supplementary Table 1**) and monomeric (119 AA) and dimeric (210 AA) versions of titin (all labeled at Cys, see **Table 1**). Plotting the total time (Δτ, see Figure 3a) that a substrate was bound and processed by ClpXP versus the length of the substrates showed a linear increase with an average processing speed of 23.9 amino acids per second (Fig. 3b and **Supplementary Fig. 2**), which agrees with previous results obtained from both bulk (28) and single-molecule assays (11, 12, 29). We obtained a similar processing speed of 14.5 amino acids per second (translocation of 16 amino acids for 1.1 seconds) from the doubly-labeled titin (**Supplementary Figure 3b**). In Figure 3b, the y-axis offset of 4.2 s reports on the initial docking phase and the eventual retention within ClpP_14_. Our data indicates that the ClpXP fingerprinter has the potential to determine both the order and spacing distance of labeled residues.

**Figure 3.**
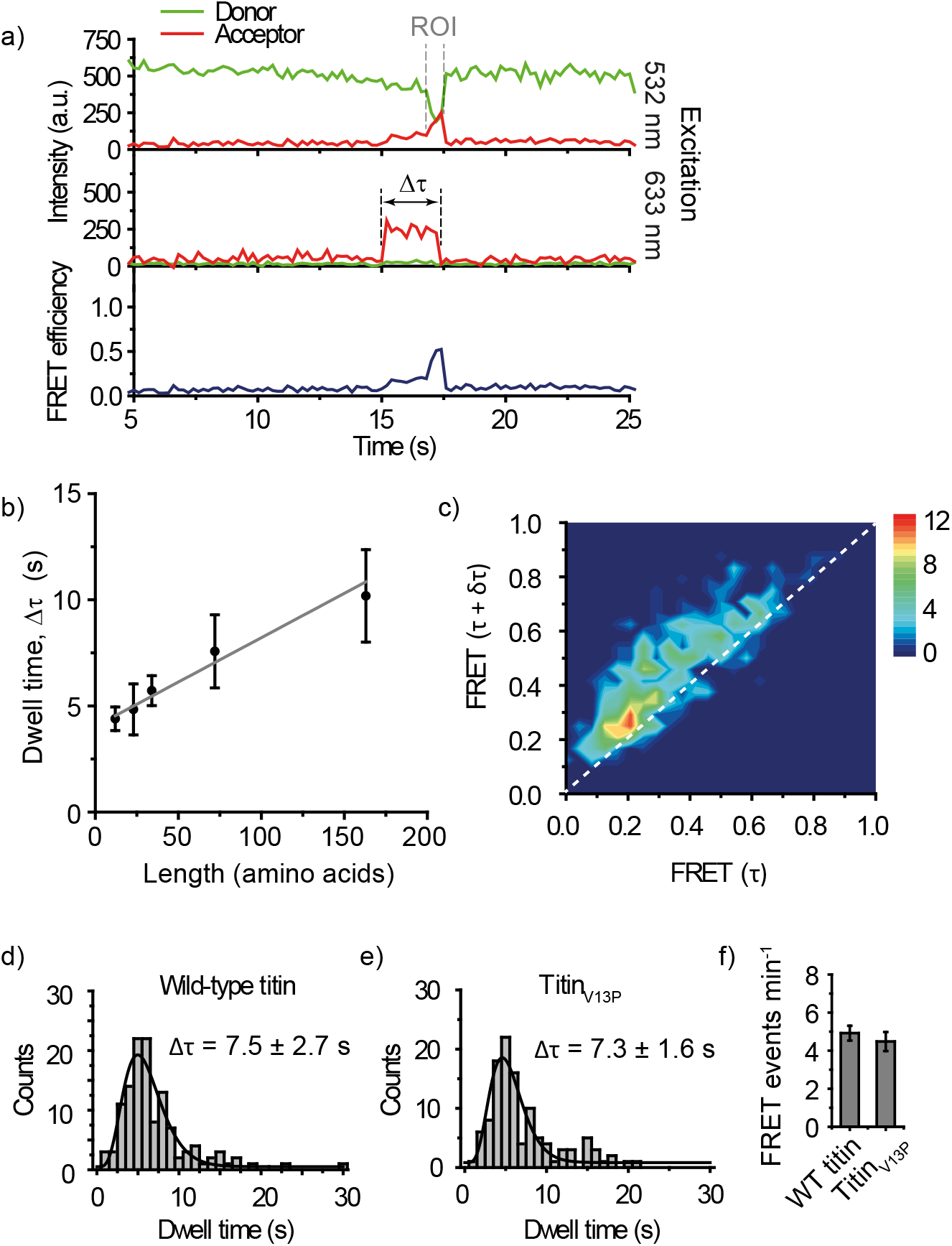
ClpXP performs uni-directional scanning with a constant speed. **(a)** Representative time trace. ROI (Region of Interest) is where the FRET efficiency gradually increases. Δτ: the total docking time. **(b)** Total dwell time vs. substrate length. The average time, <Δτ>, was obtained by fitting data in **Supplementary Figure 4** with a gamma distribution. Five different substrates were used: K-16-C-ssrA (n = 227), K-16-C-11-ssrA (n = 131), K-16-C-22-ssrA (n = 290), titin monomer (n = 85), titin dimer (n = 81). The substrate length is the number of amino acids between the C-terminus and a dye the most proximal to the N-terminus. Error bars obtained by bootstrapping with 1000 resamples. A linear fit results in an offset of 4.0 ± 0.20 s and a speed of 23.9 ± 2.86 amino acids/s. **(c)** Transition density plot. FRET change was analyzed by measuring FRET_t_ = _τ_ and FRET_t_ = _τ_ + _δτ_, with δτ = 0.4 s, for every point in ROI. The dotted line represents FRET_δτ_ = FRET_τ_ + _δτ_. K-38-C-ssrA was used. **(d)** and **(e)** Total dwell times (Δτ) for wild-type titin and titin_V13P_. Δτ = 7.5 ± 2.7 s and 7.3 ± 1.6 s were obtained respectively by fitting with a gamma distribution. Errors obtained by bootstrapping with 1000 resamples. Wild-type titin, *n* = 123. Titin_V13P_, *n* = 112. **(f)** The number of traces showing FRET events for wild-type (WT) titin and titin_V13P_. Error bars are standard deviations from 15 measurements.

Uni-directional translocation is also of utmost importance for our new technology. Backtracking of ClpXP would result in insertion errors in the observed fingerprint and thus reduce the detection precision. To evaluate the occurrence of backtracking, we determined the change in FRET over time during processing of peptide substrates. We created a 2D heat map by plotting the change of FRET over a given time interval δτ. In Figure 3c, FRET (t = τ + δτ) versus FRET (t = τ) is deposited for every time point along a time trace reporting on translocation (Figure 3a, ROI (region of interest)). We set δτ = 0.4 s, a time scale longer than our time resolution (0.2 s) but shorter than the average translocation time (0.7 s, Fig. 1d), to visualize the gradual increase of FRET. Any backtracking of ClpX_6_ along the substrate would result in momentary FRET decrease during translocation, which would appear as FRET (t = τ + δτ) values lower than FRET (t = τ) (population below the diagonal). We observed FRET (t = τ + δτ) higher than FRET (t = τ) (upper diagonal population) for a major fraction (92.5 %) of the data points. The remaining fraction is likely due to the backtracking of ClpX, the statistical noise of the fluorescence signals, and the photoblinking of acceptor dyes. This degree of experimental error is predicted not to interfere with the ability to extract length information according to our computational simulation (10).

A single-molecule protein fingerprinter should be able to process any structural element of a protein. Single-molecule force spectroscopy studies of ClpXP have shown that ClpX_6_ stalls on substrates with rigid secondary structures (30, 31), which would inhibit the extraction of sequence information. We therefore explored the possibility of disrupting such tightly folded structures to enable fingerprinting. Perturbation of cysteine residues in the titin protein has been shown to interfere with the secondary structure of the protein, making it behave as an unstructured polypeptide chain (32, 33). We purified the I27 domain of both wild-type titin, known to make ClpX_6_ stall (30), and titin_V13P_, a variant that is still folded but is degraded at a rate close to denatured titin (33). By fluorophore labeling the cysteine residues of wild-type titin and titin_V13P_, we sought to determine the degree of structural influence of the cysteine-dye conjugation on ClpXP processing. We obtained equivalent total dwell times for processing stable wild-type titin (Δτ = 7.5 ± 2.7 s, Fig. 3d) and titin_V13P_ (Δτ = 7.3 ± 1.6 s, Fig. 3e). A similar number of both substrates was processed by ClpXP within our time interval of observation (Fig. 3f), indicating that ClpX_6_ can process labeled wild-type and V13P substrates with the same efficiency. These results suggest that preparing substrates for sequencing by labeling cysteine residues (and likely lysine residues as well) might sufficiently destabilize their protein structures. This will allow for fingerprinting of any protein regardless of structural stability.

## Discussion

We have demonstrated a FRET-based detection platform utilizing an AAA+ protease as a scanner of peptides and proteins. In our approach, we conjugate fluorophores to thiol groups of cysteine residues and amine groups of the N-terminal site (which can be extended to lysine residues) because these chemical groups can be labeled with high efficiency and specificity. Our platform, however, is not limited to these two modifications. With appropriate chemistry, one could target other residues or even post-translational modifications. Detection of these moieties could be implemented by extending our current three-color FRET scheme to four-color FRET (21).

Several outstanding perspectives remain in order for our method to be directly applied to a protein sequencing technology. First, for proteomics analysis, our sequencing technique has to work for all cellular proteins without sequence bias. ClpX, a core of our platform, only recognizes substrates displaying specific sequence tags including ssrA. The substrate selectivity of ClpX would need to be broadened, perhaps through targeted mutations in the substrate-recognition loops of the ClpX channel, or the use of engineered adaptor proteins (e.g. modified SspB) that non-specifically deliver substrates to ClpX. Second, a challenge of cellular protein analysis is to detect low-abundance proteins within a complex sample such as a clinical tissue sample. The depth of the sequencing coverage might be increased by removing housekeeping proteins chromatographically (34). Third, to cover the whole proteome in a reasonable amount of time, the throughput should be enhanced. Under the standard conditions used in this work (10 nM substrate, 512x512 pixel camera, ClpX_6_), we obtained ~10 productive reactions per minute per imaging area. By using a CMOS camera that has a larger number of pixels (e.g. 2084x2084 pixels from (Juette et al, (35)) as well as a zero-mode waveguide platform that allows for single-molecule imaging of a higher concentration of substrate (e.g. 1 μM) (36), the throughput would be improved by a factor of ~1000. We also observed that productive reactions (a trace ending with high FRET) make up only ~10% of the total population (**Supplementary Figure 2d**). We suspect that this low yield is due to the lack of the N-terminal domain of ClpX in hexameric linked ClpX. By using wildtype, monomeric ClpX and also introducing an adaptor protein that facilitates substrate binding, we expect that the percentage of the productive population will reach near 100%. When our sequencer is improved with these changes, we expect to cover 1x of a single human-cell proteome (~10^8^ proteins) in nearly 10 hours (10 events /min * 16 * 100 * 10 ≈ 10^7^ events / hour).

Our method has the potential to scan full-length proteins from end to end without the need for fragmentation. Sequencing substrates are processed at a constant speed, allowing for accurate protein identification.(10) In this proof-of-concept study we show our capability to detect low-frequency subpopulations of differentially labeled substrates as well as our capacity to detect distinct acceptor fluorophores on a single substrate in a sequential manner. The platform we present here has the capability to transform proteomics from a basic research tool into an invaluable asset to clinical diagnostics.

## Methods

### ClpX_6_ purification and biotinylation

To ensure proper immobilization and hexamer formation of ClpX at low concentrations, ClpX_6_ (ΔN), a covalently linked hexamer containing a single biotinylation site, was used throughout the experiments. ClpX_6_ (ΔN) was overexpressed and purified as described.(37) In brief, ClpX_6_ protein expression and biotinylation was induced in a *E. coli* BLR(DE3) strain at O.D._600_~0.6 by adding 1.0 mM IPTG and 100 μM of biotin to increase BirA-mediated biotinylation efficiency. The culture was incubated overnight at 18°C. Cells were pelleted and resuspended in lysis buffer (20 mM HEPES pH 7.6, 400 mM NaCl, 100 mM KCl, 10% glycerol, 10 mM β-mercaptoethanol, 10 mM imidazole) in the presence of 1 mM PMSF and lysed by French press twice at 20 psi. ClpX_6_(ΔN) was purified from the supernatant first with Ni^2+^-NTA affinity resin, followed by size exclusion chromatography with a Prep Sephacryl S-300 16/60 High Resolution column (GE Healthcare).

### ClpP mutations, purification and labeling

Point mutations were constructed in ClpP by overlap extension PCR to produce the cysteine-free variant ClpP_C91S-C113S_, and the subsequent variants ClpP_Q48C_, ClpP_A139C_ and ClpP_F31C_. The variants were overexpressed in *E. coli* BL21(DE3)pLysS at O.D._600_~0.6 by adding 0.5 mM IPTG and incubated for 3 h at 30°C. Cells were pelleted and resuspended in lysis buffer (50 mM sodium phosphate pH 8.0, 1 M NaCl, 10% glycerol, 5 mM imidazole) in the presence of Set III protease inhibitors (Calbiochem) and lysed by French press twice at 20 psi. ClpP was purified from the supernatant first with Ni^2+^-NTA affinity resin, followed by size exclusion chromatography with a Prep Sephacryl S-300 16/60 High Resolution column (GE Healthcare). ClpP was dialyzed overnight against PBS (pH 7.4) before labeling for 4 h at 4°C with monoreactive maleimide donor dye (Cy3, GE Healthcare, for two-color experiments, and Alexa488, Invitrogen, for three-color experiments). 10x molar dye excess was used in PBS pH 7.4 under nitrogen. Free dye was removed using PD Minitrap G-25 size exclusion columns (GE Healthcare). Labeling efficiency of 5.9, 1.1, and 1.7 dyes per tetradecameric ClpPQ_48C_, ClpP_A139C_, and ClpP_F31C_, respectively, was measured by spectrophotometry (DeNovix DS-11 FX).

### ClpP inactivation

Purified ClpPQ48C was chemically inactivated as described previously (1). Briefly, ClpPQ48C (4 μM) was inactivated in PD buffer containing 10 mM DFP (Sigma). The reaction was incubated for 6 h at 4°C and then dialyzed twice: 1x 2 h and 1x overnight against 1x PBS (pH 7.4). ClpPQ_48C_DFP_ was labeled with monoreactive maleimide donor dye, Cy3, for 4 hours at 4 °C. A 10x molar excess of dye was used in PBS pH 7.4 under nitrogen. Free dye was removed using Pierce^™^ Dye Removal Columns (Thermo Fisher). A labeling efficiency of 1.8 dye per tetradecameric ClpPQ_48C_DFP_ was measured by spectrophotometry (DeNovix DS-11 FX).

### ClpXP cleavage reaction

To assess the enzymatic activity of donor-labeled ClpXP, 0.9 μM ClpX and 2.9 μM of ClpP (WT or variants) in PD buffer (25 mM HEPES pH 8.0, 5 mM MgCl_2_, 40 mM KCl, 0.148% NP-40, 10% glycerol) were incubated at 30°C in the presence of 10 μM titin_V13P_-ssrA and 5 mM ATP. Samples were taken at t = 0 min and 30 min and analyzed using 4 – 20% precast SDS-PAGE gels (Thermo Scientific) and coomassie staining.

### Substrate preparation

Titin-I27 (wild-type, V13P and dimer) with the C-terminal ssrA tag was expressed in *E. coli* BL21AI at 0.D._600_~0.6 by adding 0.2 % arabinose and incubated for 4 h at 37°C. Cells were pelleted and resuspended in lysis buffer (50 mM sodium phosphate pH 8.0, 500 mM NaCl, 10 mM imidazole), then lysed by sonication. Titin was purified from the supernatant with Ni^2^+-NTA affinity resin. Titin was dialyzed overnight against PBS (pH 7.4) before labeling for 4 h at 4°C with 10× molar excess of monoreactive maleimide acceptor dye (Cy5, GE Healthcare) in the presence of 4 M GdnCl in PBS pH 7.4 under nitrogen. Custom designed polypeptides were obtained from Biomatik. Cysteine residues of the polypeptides were labeled with monoreactive maleimide-functionalized Cy5 as an acceptor for two-color measurements and with Cy3 and Cy5 as an acceptor for three-color measurements. Polypeptides were labeled in the presence of a 10x molar excess of dye overnight at 4°C in PBS under nitrogen. For labeling with additional acceptors at N-terminus, monoreactive NHS-ester functionalized dyes (Cy3 or Cy5, GE Healthcare) were added to the reaction mixture described above, also in 10x molar excess. Free dye was removed using PD Minitrap G-25 size exclusion columns (GE Healthcare). Labeling efficiencies up to 95% were measured by spectrophotometry (DeNovix DS-11 FX) (See Supplementary Table 1 for the full list of substrates.)

### Single-molecule sample preparation

To reduce the nonspecific binding of proteins, acidic piranha-etched quartz slides (G. Finkenbeiner) were passivated with two rounds of polyethylene glycol (mPEG-Succinimidyl Valerate, MW 5000 Laysan, followed by MS(PEG)_4_, Piercenet) as described previously (38). After assembly of a microfluidic flow chamber, slides were incubated with 5% Tween-20 for 10 min (39), and excess Tween-20 was washed with T50 buffer (10 mM Tris-HCl pH 8.0, 50 mM NaCl), followed by 1 minute incubation with streptavidin (0.1 mg/ml, Sigma). Unbound streptavidin was washed with 100 μL of T50 buffer, followed by 100 μL of PD buffer (25 mM HEPES pH 8.0, 5 mM MgCl_2_, 40 mM KCl, 0.148 % NP-40, 10 % glycerol). A ClpX_6_:ClpP_14_ = 1:3 molar ratio was used to ensure ClpXP complex formation with a 1:1 molar ratio (40). 30 nM ClpX_6_ and 90 nM ClpP_14_ (either wild-type or mutant) were preincubated for 2 min at room temperature in the presence of 10 mM ATP in PD buffer. After preincubation, the sample was diluted 10 times in PD buffer to reach an expected final ClpXP complex concentration of 3 nM. The diluted sample was applied to the flow chamber and incubated for 1 min. Unbound ClpXP complexes were washed with 100 μL PD buffer containing 1 mM ATP. 10 – 20 nM of acceptor-labeled substrate was introduced to the flow chamber in the presence of an imaging buffer (0.8% dextrose (Sigma), 1 mg/mL glucose oxidase (Sigma), 170 mg/mL catalase (Merck), and 1 mM Trolox ((±)-6-Hydroxy-2,5,7,8-tetramethylchromane-2-carboxylic acid, 238813), (Sigma)). Donor-labeled ClpP_14_ added into a chamber without ClpX_6_ led to very few non-specifically immobilized ClpP protein complexes, ruling out any nonspecific adsorption of ClpP_14_ to the surface (**Supplementary Fig. S1b**). All experiments were performed at room temperature (23 ± 2°C).

### Single-molecule fluorescence

Single-molecule fluorescence measurements were performed with a prism-type total internal reflection fluorescence microscope. For two-color measurements, Cy3 molecules were excited using a 532 nm laser (Compass 215M-50, Coherent), and Cy5 molecules were excited using a 633 nm laser (25 LHP 928, CVI Melles Griot). Fluorescence signals from single molecules were collected through a 60x water immersion objective (UplanSApo, Olympus) with an inverted microscope (IX71, Olympus). Scattered light from the 532 nm and 633 nm laser beams was blocked by a triple notch filter (NF01-488/532/635, Semrock). The Cy3 and Cy5 signals were separated with a dichroic mirror (635 dcxr, Chroma) and imaged using an EM-CCD camera (Andor iXon 897 Classic, Andor Technology).

For three-color measurements, Alexa488 molecules were excited using a 473 nm laser (OBIS 473 nm LX 75 mW, Coherent), Cy3 molecules were excited using a 532 nm laser (Sapphire 532nm-100 CW, Coherent), and Cy5 molecules were excited using a 637 nm laser (OBIS 637 nm LX 140 mW, Coherent). Fluorescence signals from single molecules were collected through a 60x water immersion objective (UplanSApo, Olympus) with an inverted microscope (IX73, Olympus). The 473 nm laser beam was blocked by a 473 nm long pass filter (BLP01-473R-25, Semrock), the 532 nm laser beam was blocked by a 532 nm notch filter (NF03-532E-25, Semrock), and the 637 nm laser beam was blocked by a 633 nm notch filter (NF03-633E-25, Semrock). The Alexa488, Cy3 and Cy5 signals were separated by dichroic mirrors (540dcxr and 635 dcxr, Chroma) and imaged using an EM-CCD camera (Andor iXon 897 Classic, Andor Technology).

### Data acquisition

Samples were excited alternatingly with different colors and using a custom-made program written in Visual C++ (Microsoft). A series of CCD images with an exposure time of 0.1 s was recorded. The time traces were extracted from the CCD image series using an IDL (ITT Visual Information Solution) algorithm that identifies fluorescence spots with a defined Gaussian profile and with signals above the average of the background signals. Colocalization between Alexa488, Cy3 and Cy5 signals was carried out with a custom-made mapping algorithm written in IDL. The extracted time traces were processed using Matlab (MathWorks) and Origin (Origin Lab).

## Contributions

C.J. conceived the project. J.v.G., A.S.M., and C.J. designed the research. J.v.G. and M.F. performed the single-molecule experiments. J.v.G., M.F., M.S. and P.T. purified and labeled peptides and proteins. J.v.G., and M.F. analyzed the data. J.v.G., M.F., A.S.M. and C.J. discussed the data. J.v.G., M.S., A.S.M. and C.J. wrote the manuscript.

## Acknowledgements

We would like to thank Marek Noga, Ivo Severins and Anna Haagsma for technical support; Stanley Chandradoss and Margreet Docter for building the multi-color optical setup; Luuk Loeff and Misha Klein for assisting with data analysis; and Tim Blosser, Laura Restrepo, Margreet Docter, Stephanie Heerema, Noortje de Haan, and Viktorija Globyte for critical reading of the manuscript. ClpX_6_ expression plasmids were a generous gift from Andreas Martin. ClpP expression plasmids were a generous gift from Tania Baker and Robert Sauer. Titin expression plasmids were a generous gift from Victor Muñoz and Jörg Schönfelder. C.J. and A.S.M were funded by the Foundation for Fundamental Research on Matter (12PR3029).

